# On the Geometry of Elementary Flux Modes

**DOI:** 10.1101/2022.09.24.509324

**Authors:** Frederik Wieder, Martin Henk, Alexander Bockmayr

## Abstract

Elementary flux modes (EFMs) play a prominent role in the constraint-based analysis of metabolic networks. They correspond to minimal functional units of the metabolic network at steady-state and as such have been studied for almost 30 years. The set of all EFMs in a metabolic network tends to be very large and may have exponential size in the number of reactions. Hence, there is a need to elucidate the structure of this set. Here we focus on geometric properties of EFMs. We analyze the distribution of EFMs in the face lattice of the steady-state flux cone of the metabolic network and show that EFMs in the relative interior of the cone occur only in very special cases. As a measure of complexity, we introduce the concept of the degree of an EFM, which is the dimension of the inclusionwise minimal face containing it. Geometric analysis can help to better understand the structure of the set of EFMs, which is important from both the mathematical and the biological viewpoint.

## 1 Introduction

Constraint-based analysis of metabolic networks is an important area in computational biology [2, 3]. The stoichiometric and thermodynamic constraints that have to hold in a metabolic network at steady-state have led to the definition of the *steady-state flux cone*, which comprises all possible flux distributions over the network at steady-state.

An important concept to analyze the flux cone in a mathematically and biologically meaningful way are *elementary flux modes* (EFMs) [15, 16], which provide an inner description of the flux cone based on generating vectors [4, 19, 9, 5]. From a mathematical point of view, not all elementary flux modes are needed to describe the cone, i.e., the set of all EFMs does not form a minimal generating set, except for special cases (for example when all reactions are irreversible). Even for small networks, the number of elementary flux modes may be very large.

[10] introduced metabolic behaviors and studied outer descriptions of the flux cone based on *minimal metabolic behaviors* (MMBs), which are in a one-to-one correspondence with the minimal proper faces of the flux cone. [12] introduced the concept of a minimal set of elementary flux modes (MEMo) and gave an algorithm to compute such a set. A MEMo consists of an EFM from each minimal proper face of the flux cone together with a vector space basis of the lineality space. In general, a minimal proper face may contain many different EFMs.

The goal of this paper is to get a deeper understanding of the structure of the set of EFMs by further studying their geometric properties. We generalize the result of [10] and show that higher-dimensional faces of the flux cone can be characterized by metabolic behaviors. We study the distribution of EFMs over the faces of the flux cone and introduce as a measure of complexity the *degree* of a flux vector or EFM, which is the dimension of the inclusionwise minimal face containing it. We prove upper bounds on the degree of EFMs and show that EFMs in the relative interior of the flux cone occur only in very special cases. We also study the effect of removing blocked reactions and redundant irreversibility constraints and deduce an upper bound on the cardinality of minimal metabolic behaviors.

## 2 Polyhedral cones

In this section we introduce some mathematical background on polyhedral cones. For further reading we refer to [11, 13, 14].

A vector *x* ∈ ℝ^*n*^ is a *linear combination* of the vectors *x*^1^,…,*x^k^* ∈ ℝ^*n*^ if *x* = λ_1_*x*^1^+⋯+λ_*k*_ *x^k^*, for some λ_1_,…, λ_*k*_ ∈ ℝ. If, in addition

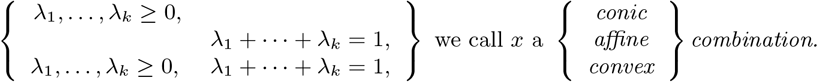

For a nonempty subset *X* ⊆ ℝ^*n*^, we denote by lin(*X*) (resp. cone(*X*), aff(*X*), conv(*X*)) the *linear* (resp. *conic, affine, convex*) *hull of X*, i.e., the set of all linear (resp. conic, affine, convex) combinations of finitely many vectors of *X*.

A nonempty set *C* ⊆ ℝ^*n*^ is a *convex cone*, if it is closed under conic combinations, i.e., λ_*x*_ + *μy* ∈ *C*, for all *x,y* ∈ *C* and *λ,μ* ≥ 0. A convex cone *C* is *polyhedral* if it is the solution set of a system of finitely many homogeneous linear inequalities, i.e., if

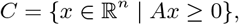

for some matrix *A* ∈ ℝ^*m*×*n*^. If a cone C is the conic hull of a finite set *X* = {*x*^1^,…,*x^s^*} ⊂ ℝ^*n*^, it is called *finitely generated* and the set *X* is called a *generating set* of *C*. By the well-known theorem of Farkas-Minkowski-Weyl (see e.g. [14]), a convex cone is polyhedral if and only if it is finitely generated. For the rest of this paper we will only consider polyhedral cones and often simply write cone.

If *C* = {*x* ∈ ℝ^*n*^ | *Ax* ≥ 0}, is a polyhedral cone, an inequality *ax* ≥ 0, where *a* denotes a row of A and *ax* the inner product of *a* and *x*, is called an *implicit equality in Ax* ≥ 0, if *ax* = 0, for all *x* ∈ *C*. Following [14], we denote by *A*^=^*x* ≥ 0 the system of implicit equalities in *Ax* ≥ 0 and by *A*^+^*x* ≥ 0 the remaining inequalities.

If removing an inequality *ax* ≥ 0 from *Ax* ≥ 0 does not change the associated cone *C*, the inequality is called *redundant*. If there are no redundant inequalities, the description *Ax* ≥ 0 is called *irredundant*.

The *dimension* dim(*C*) of a cone *C* is the dimension of its affine hull aff(*C*) = {*x* ∈ ℝ^*n*^ | *A*^=^*x* = 0} and is equal to *n* – rank(*A*^=^). Since 0 ∈ *C*, aff(*C*) coincides with the linear hull lin(*C*).

A vector *x* ∈ *C* is in the *relative interior of C*, shortly *x* ∈ relint(*C*), if there exists *ϵ* > 0 such that *B_ϵ_*(*x*) ⋂ aff(*C*) ⊆ *C*, where *B_ϵ_*(*x*) is the *n*-dimensional ball of radius *ϵ* centered at *x*. If *x* ∈ *C* is not in the relative interior of *C*, it is in the *relative boundary of C*.

The *lineality space* of a cone *C* = {*x* ∈ ℝ^*n*^ | *Ax* ≥ 0} is given by lin. space(*C*) = {*x* ∈ ℝ^*n*^ | *Ax* = 0}, which is the inclusionwise maximal linear subspace contained in *C*. A cone *C* is called *pointed* if its lineality space is trivial, i.e., lin. space(*C*) = {0}. If a cone is pointed, it does not contain a line.

An inequality *ax* ≥ 0 is called *valid* for *C* if *C* ⊆ {*x* ∈ ℝ^*n*^ | *ax* ≥ 0}. A nonempty set *F* ⊆ *C* is called a *face* of *C* if there exists an inequality *ax* ≥ 0 valid for *C* such that *F* = *C* ⋂ {*x* ∈ *C* | *ax* = 0}. The hyperplane {*x* ∈ *C* | *ax* = 0} is then called a *supporting hyperplane* of *F*. Alternatively, a face can be characterized as a nonempty set *F* ⊆ *C* with *F* = {*x* ∈ *C* | *A_I*_x* = 0}, where *A_I_,\** is the submatrix of *A* whose rows belong to the set *I* ⊆ {1,…,*m*} [14].

A polyhedral cone *C* has only finitely many faces, each face *F* of *C* is itself a polyhedral cone and *F*′ ⊆ *F* is a face of F if and only if *F*′ is a face of *C*. A *k*-dimensional face will also be called a *k-face*. A cone *C* is pointed if and only if it has a 0-face, namely the origin.

A face *F* ≠ *C* of *C* is called a *facet* if it is inclusionwise maximal, i.e., there is no other face *F*′ = *C* such that *F* ⊂ *F*′. If the description *Ax* ≥ 0 of *C* is irredundant, there is a 1-1 correspondence between the facets of *C* and the inequalities in *A*^+^*x* ≥ 0 [14, Theor. 8.1]. In particular, for every facet *F* there is an inequality *ax* ≥ 0 from *A*^+^*x* ≥ 0 such that *F* = {*x* ∈ *C* | *ax* = 0}. We have dim(*F*) = dim(*C*) – 1 for every facet *F* of *C*, and every face of *C* (except *C* itself) is the intersection of facets of *C*.

The faces of a polyhedral cone form a lattice under set inclusion, called the *face lattice*. If two polyhedral cones have isomorphic face lattices, they are called *combinatorially equivalent*.

#### Proposition 2.1.

*Let C* = {*x* ∈ ℝ^*n*^ | *Ax* ≥ 0} *be a polyhedral cone and z* ∈ *C. Let* 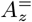 *be the submatrix of A whose rows correspond to the inequalities in Ax* ≥ 0 *that are fulfilled with equality by z. Let F be the face of C defined by* 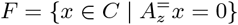. *Then*

i. *F is the inclusionwise minimal face of C containing z*,
ii. 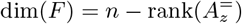,
iii. *z* ∈ relint(*F*).

*Proof.* For *x* ∈ ℝ^*n*^, define I(*x*) = {*i* ∈ {1,…,*m*} | *_i_x* = 0}, where *a_i_* is the *i*-th row in *A*. Let *F*′ = {*x* ∈ *C* | *A*_*I,\**_*x* = 0} be a face of C containing *z*. Then *A*_*I*,*_ *z* = 0 and thus *I* ⊆ *I*(*z*), which implies *F* = {*x* ∈ *C* | *A*_*I*(*z*),*_ *x* = 0} ⊆ *F*′ and statement i) follows.

For *x* ∈ *F*, we have *I*(*z*) ⊆ *I*(*x*). Therefore, *I*(*z*) has the minimal number of elements among *I*(*x*), where *x* ∈ *F*. The statements ii) and iii) now follow from Prop. 4.3 in [11] and its proof.

If *C* is a cone with dim(lin. space(*C*)) = *t* ≥ 0, a face of dimension *t* + 1 is called a *minimal proper face* of *C*. For a pointed cone *C*, the minimal proper faces are the 1-faces, which are called the *extreme rays* of *C*. Equivalently, cone({*r*}) ⊆ *C*, *r* = 0, is an extreme ray of *C* if and only if *r* = *x* + *y* implies *x, y* ∈ cone({*r*}), for all *x, y* ∈ *C*.

The *Minkowski sum* of two sets *X* and *Y* is defined as *X* + *Y* = {*x* + *y* | *x* ∈ *X, y* ∈ *Y*}. The next result states that any polyhedral cone can be decomposed into a Minkowski sum of its lineality space and a pointed cone.

#### Proposition 2.2.

*Let C* ⊆ ℝ^*n*^ *be a polyhedral cone, L* = lin. space(*C*). *Let G*^1^,…,*G^s^,s* ≥ 0, *be the distinct minimal proper faces of C and ^i^* ∈ *G^i^* \ *L, for i* = 1,…, *s. Let P* = cone({*g*^1^,…,*g^s^*}). *Then*

i. *P is a pointed cone and its extreme rays are* cone({*g*^1^}),…,cone({*g^s^*}),
ii. *C* = *L* + *P* = *L* + cone({*g*^1^,…, *g^s^*}), *L* ⋂ *P* = {0} *and if L* ⋂ lin(*P*) = {0} *then* dim(*C*) = dim(*L*) + dim(*P*).

*Proof.* i) By definition *P* is a finitely generated cone. Assume that *P* is not pointed. Then there exist λ_*i*_ ≥ 0, i = 1,…, *s*, not all equal to zero, such that 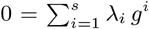. Hence, there exists *j* ∈ {1,…, *s*} such that –*g^j^* ∈ *P* ⊆ *C* and so *g^j^* ∈ *L*, contradicting our choice.

To see that *g*^1^,…,*g^s^* define extreme rays of *P*, assume without loss of generality that cone({*g*^1^}) is not an extreme ray of *P*. Then we can find *μ_i_* ≥ 0, 2 ≤ *i* ≤ *s*, not all equal to zero, such that 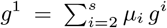. As *G*^1^ is a face, there exists a supporting hyperplane *H_a_* := {*x* ∈ ℝ^*n*^ | α*x* = 0}, *a* ∈ ℝ^*n*^ \ {0}, such that *G*^1^ = *H_a_* ⋂ *C* and α*x* > 0 for all *x* ∈ *C* \ *G*^1^. Thus, from 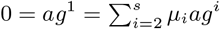 we conclude that *ag^k^* = 0 for some *k* ∈ {2,…,*s*}. Therefore *g^k^* ∈ *G*^1^ and *G^k^* ⊆ *G*^1^, which leads to *G^k^* = *G*^1^, because *G*^1^ is a minimal proper face. But then *G*^1^,…,*G^s^* are not distinct, which is a contradiction.

ii) By Theorem 8.5 in [14], we have *C* = *L*+*P*. Since *L* is a face of *C*, there exists *a* ∈ ≥” \{0} such that *ax* = 0 for all *x* ∈ *L* and *ax* > 0 for all *x* ∈ *C* \ *L*. From *ag^i^* > 0, *i* = 1,…, *s*, we get *ax* > 0, for all *x* ∈ *P* \ {0}, hence *L* ⋂ *P* = {0}. If *L* ⋂ lin(*P*) = {0}, we have lin(*C*) = *L* ⊕ lin(*P*), where *L* ⊕ lin(*P*) is the direct sum of the vector spaces *L* and lin(*P*), and so dim(*C*) = dim(*L*) + dim(*P*).

The combinatorial type of the pointed cone *P* = cone({*g*^1^,…, *g^s^*}) in Prop. 2.2 is (in general) not uniquely determined. However, if we choose *g*^1^,…,*g^s^* such that *L* ⋂ lin(*P*) = {0}, i.e., all the *g^i^* are contained in some linear subspace *L*′ complementary to *L*, then the combinatorial type of *P* is independent of the choice of the *g^i^* from the minimal proper faces. Observe that for any complementary space *L*′ of *L, L*′ ⋂ *G^i^* is a ray, i.e., *L*′ ⋂ *G^i^* = cone({*g^i^*}) for some *g^i^* ∈ *L*′ ⋂ *G^i^*.

#### Proposition 2.3.

*Let C* ⊆ ℝ^*n*^ *be a polyhedral cone, L* = lin. space(*C*), *and let P*_1_, *P*_2_ *be pointed cones with L* + *P*_1_ = *C* = *L* + *P*_2_ *and L* ⋂ lin(*P*_1_) = {0} = *L* ⋂ lin(*P*_2_). *Then P*_1_ *and P*_2_ *are combinatorially equivalent*.

*Proof.* Without loos of generality let dim(*C*) = *n* and dim(*L*) = *t*. With 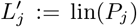, by Prop. 2.2 we have 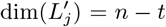 and 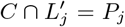, for *j* = 1, 2. Let *u*^1^,…, *u^n–t^* be a basis of 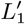. As also 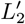 is a complement of *L*, there exist uniquely determined 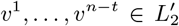, *w*^1^,…, *w^n–t^* ∈ *L* such that *u^i^* = *v^i^* + *w^i^*,1 ≤ *i* ≤ *n* – *t*. Now let *T* : ℝ^*n*^ → ℝ” be the invertible linear map with

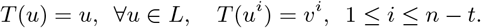

Then we get *T*(*C*) = *C*. To see this let *y* = *u* + *w* ∈ *C* with 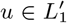 and *w* ∈ *L*. Then we may write

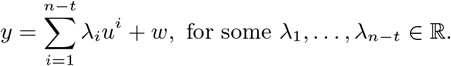

Thus

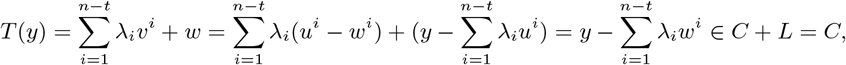

and vice versa. We conclude

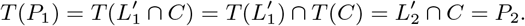

Thus, *P*_1_ and *P*_2_ are affinely and thereby also combinatorially equivalent.

We also point out that the relative interior of a polyhedral cone can easily be described by looking at the implicit equalities in *Ax* ≥ 0:

#### Proposition 2.4.

*Let C* = {*x* ∈ ℝ” | *Ax* ≥ 0} = {*x* ∈ ℝ^*n*^ | *A*^=^*x* = 0, *A*^+^*x* ≥ 0} *be a polyhedral cone. Then*

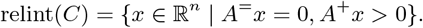

*Proof.* If *x* ∈ *C* with *A*^+^*x* > 0 then for any *y* ∈ lin(*C*) = {*y* ∈ ℝ^*n*^ | *A*=*y* = 0} there exists *ϵ* > 0 such that *A*^+^(*x* + *ϵy*) > 0. Hence *x* ∈ relint(*C*). Conversely, let *x* ∈ relint(*C*) and let *a* be an arbitrary row of *A*^+^. By definition of *A*^+^ there exists *z* ∈ *C* with *az* > 0. As *x* ∈ relint(*C*), there exists *ϵ* > 0 such *x* – *ϵz* ∈ *C* and so *a*(*x* – *ϵz*) ≥ 0. Thus *ax* > 0.

## 3 Metabolic networks

A *metabolic network* 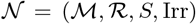 is given by a set 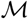 of (internal) *metabolites*, a set 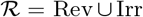 of *reversible* and *irreversible reactions*, and a *stoichiometric matrix S* ∈ ℝ^*m*×*n*^, where 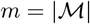 and 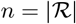. For 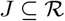, we denote by *S_*,J_* the submatrix of *S* whose columns belong to *J*.

The network can be seen as a weighted hypergraph with the generalized incidence matrix *S*, where the metabolites are represented by nodes and the reactions by hyperarcs. A positive entry *S_i,j_* > 0 in the stoichiometric matrix *S* indicates that reaction *j* produces metabolite *i*. If *S_i,j_* < 0, metabolite i is consumed in reaction *j*.

#### Example 3.1.

*The metabolic network in Fig. 1 has the set of metabolites* 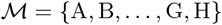, *the set of reversible reactions* Rev = {1, 3, 4, 5, 9, 10, 11, 12} *and the set of irreversible reactions* Irr = {2, 6, 7, 8, 13}. *The stoichiometric matrix is*

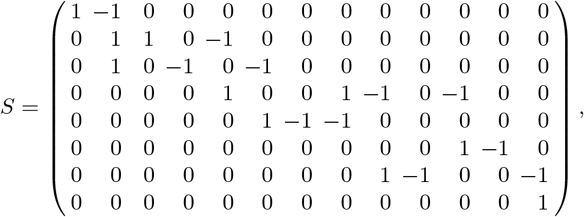

*where we assume that reversible reactions are oriented from left to right and from top to bottom.*

**Figure 1:**
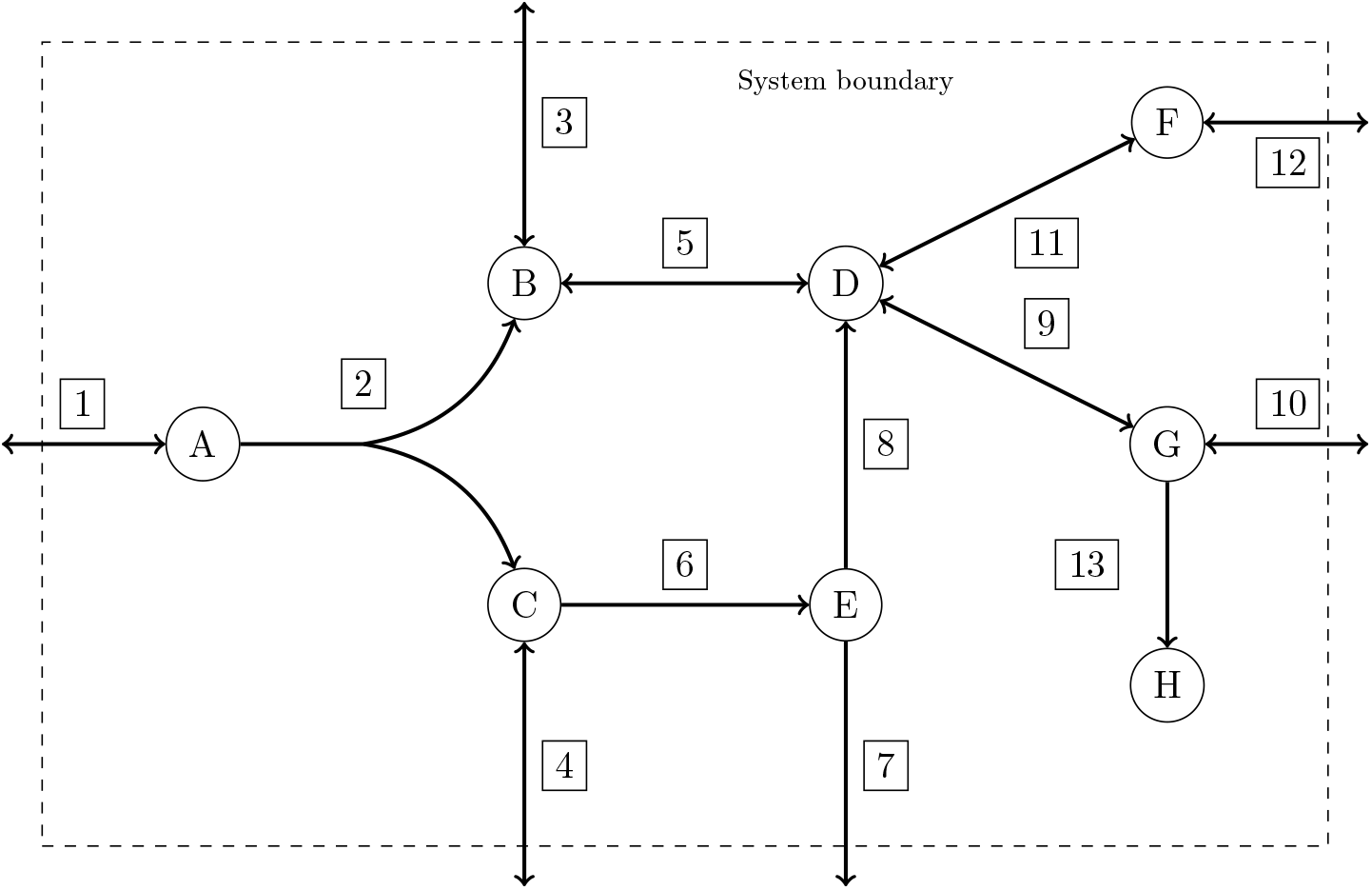
Example of a metabolic network

### 3.1 Flux cones

In metabolic network analysis, we often assume that the network is in *steady-state*, i.e., for each internal metabolite the rate of production is equal to the rate of consumption. In matrix notation, the steady-state constraints can be written as *Sv* = 0, where *v* ∈ ℝ^*n*^ denotes a *flux vector.* By adding the *thermodynamic irreversibility* constraints *v_j_* ≥ 0, for all *j* ∈ Irr, and setting

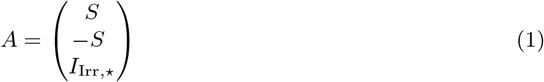

we obtain the polyhedral cone

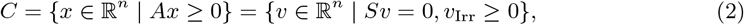

which is called the (steady-state) *flux cone* of 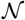. Here, *v*_Irr_ is the subvector of *v* whose components belong to Irr, and *I*_Irr,*_ the submatrix of the (*n* × *n*) identity matrix *I_n_* whose rows correspond to the irreversible reactions.

#### Implicit equalities

The implicit equalities in the definition (2) of a flux cone *C* include all steady-state constraints *Sv* = 0. If any of the irreversibility constraints *v_j_* ≥ 0, *j* ∈ Irr, is an implicit equality, the corresponding reaction *j* ∈ Irr is *blocked*, i.e., *v_j_* = 0, for all *v* ∈ *C*. For some of our results, we will assume that there are no implicit equalities in *v*_Irr_ ≥ 0, or equivalently that there are no blocked irreversible reactions. Blocked reactions can be determined by solving a linear optimization problem.

#### Redundant inequalities

If in (2) one of the irreversibility constraints *v_j_* ≥ 0, *j* ∈ Irr, is redundant, the corresponding reaction *j* can be shifted from the set Irr of irreversible reactions to the set Rev of reversible reactions, without changing the flux cone *C*. The constraint *v_j_* ≥ 0 is then implied by the remaining constraints.

#### Example 3.2.

*In the metabolic network of Fig. 1, the irreversible reaction* 13 *is blocked, i.e., v*_13_ = 0, *for all v* ∈ *C, because there is no reaction to consume metabolite* H.

*The irreversibility constraint v*_6_ ≥ 0 *is redundant. There can be no flux from metabolite* E *to metabolite C because there is no reaction producing* E.

*The reversible reaction* 1 *cannot carry flux from right to left. Reaction* 1 *could be added to the set* Irr *of irreversible reactions, but then the inequality v*_1_ ≦ 0 *would be redundant.*

If redundant inequalities are removed from the description of a flux cone, the resulting irredundant description is in general not unique, because it depends on the order in which the redundant constraints are removed.

#### Proposition 3.3.

*Let C* = {*v* ∈ ℝ^*n*^ | *Sv* = 0, *v*_Irr_ ≥ 0} *be a flux cone such that none of the inequalities v_j_* ≥ 0, *j* ∈ Irr, *is redundant or an implicit equality. Then C has exactly* | Irr| *facets and each facet F has the representation*

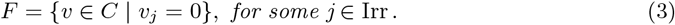

*Proof.* If there are no implicit equalities in *v*_Irr_ ≥ 0 and *A* is given by (1), then 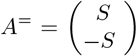 and *A*^+^ = *I*_rr,*_. Since there are no redundant inequalities in *v*_Irr_ ≥ 0, the result follows from Theorem 8.1 in [14].

### 3.2 Elementary flux modes

In metabolic network analysis we are particularly interested in minimal functional units of the network. Let *C* be the flux cone of a metabolic network 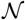. A vector *ϵ* ∈ *C* \ {0} is called an *elementary flux mode* (EFM) [15] if it has inclusionwise minimal support, i.e., if

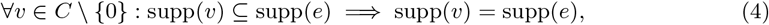

where the *support* of *v* ∈ ℝ^*n*^ is defined by 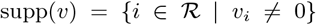. We say that a reaction 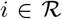 is *active* in *v* ∈ *C* if *i* ∈ supp(*v*). By irr. supp(*v*) := supp(*v*) ⋂ Irr we denote the *irreversible support* of *v*, i.e., the set of active irreversible reactions in *v*. Analogously, rev. supp(*v*) := supp(*v*) ⋂ Rev denotes the *reversible support* of *v*.

To verify whether a given flux vector *v* ∈ *C* is an EFM the *rank test* [6, 18] can be applied, i.e.,

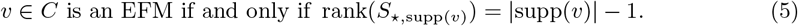

#### Proposition 3.4.

*Let C* = {*v* ∈ ℝ^*n*^ | *Sv* = 0, *v*_Irr_ ≥ 0} *be the flux cone of a metabolic network. Then*

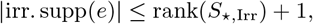

*for each EFM e* ∈ *C*.

*Proof.* Suppose the opposite. Then

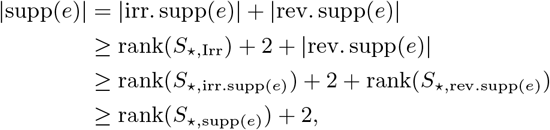

contradicting the rank test (5).

### 3.3 Illustrative examples

To illustrate the theoretical results in the following sections through concrete examples, we will use the metabolic networks `Pyruvate` and `Pentose Phosphate Pathway` from the KEGG database (https://www.genome.jp/kegg/pathway.html, [7]) and Escherichia coli str. `K-12 substr`. `MG1655` (`E.coli core`) from the BiGG database [8], where we removed the biomass reaction. The characteristics of these networks are summarized in Tab. 1. The EFMs were computed with `efmtool` (https://csb.ethz.ch/tools/software/efmtool.html, [17]). For polyhedral computations we used polymake (https://polymake.org/, [1]).

**Table 1:** Characteristics of the three example networks. 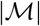 and 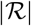 denote the number of metabolites resp. reactions, which correspond to the number of rows resp. columns of the stoichiometric matrix. |Irr| and |Rev| denote the number of irreversible resp. reversible reactions of the network. rank(*S*) is the rank of the stoichiometric matrix. The flux cone *C* = *L* + *P* is the Minkowski sum of the lineality space *L* and a pointed cone *P*, with dim(*C*) = dim(*L*) + dim(*P*). |Facets| is the number of facets of the flux cone, which is equal to the number of irreversibility constraints if none of these is redundant or an implicit equality. |blocked irr| resp. |blocked rev| describe the number of blocked irreversible resp. blocked reversible reactions. |EFMs| is the number of EFMs and | MMBs| the number of minimal metabolic behaviors (cf. Sect. 6)

|  | E. coli core | Pentose Phosphate | Pyruvate |
| --- | --- | --- | --- |
| $(m, n) = \vert \mathcal{M}\vert , \vert \mathcal{R}\vert $ | (72, 94) | (34, 57) | (28, 81) |
| $\vert \text{Irr}\vert $ | 48 | 19 | 40 |
| $\vert \text{Rev}\vert $ | 46 | 38 | 41 |
| $\text{rank}(S)$ | 67 | 34 | 28 |
| $n - \text{rank}(S)$ | 27 | 23 | 53 |
| $\dim(C)$ | 23 | 23 | 53 |
| $t = \dim(L)$ | 0 | 8 | 16 |
| $\dim(P)$ | 23 | 15 | 37 |
| $\vert \text{Facets}\vert $ | 39 | 17 | 37 |
| $\vert \text{blocked irr}\vert $ | 8 | 0 | 0 |
| $\vert \text{blocked rev}\vert $ | 2 | 0 | 0 |
| $\vert \text{EFMs}\vert $ | 16673 | 5180 | 47854 |
| $\vert \text{MMBs}\vert $ | 1421 | 19 | 37 |

## 4 Faces of the flux cone and metabolic behaviors

Given a metabolic network 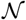 with flux cone *C*, a *metabolic behavior* [10] of *C* is a nonempty set of irreversible reactions *D* ⊆ Irr with *D* = irr. supp(*v*), for some *v* ∈ *C*. A *minimal metabolic behavior* (MMB) is a metabolic behavior *D* for which there is no other metabolic behavior 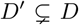. [10] have shown that minimal metabolic behaviors are in a 1-1 correspondence with the minimal proper faces of the flux cone *C*. In particular, if *G* is a minimal proper face and *L* the lineality space of *C*, then all flux vectors *v* ∈ *G* \ *L* have the same irreversible support *D_G_* = irr. supp(*v*), which is a minimal metabolic behavior.

#### Proposition 4.1.

*Let C be the flux cone of a metabolic network. Then each metabolic behavior is the union of MMBs.*

*Proof.* Let 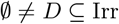 be a metabolic behavior and let *v* ∈ *C* with *D* = irr. supp(*v*). Let *L* be the lineality space and *G*^1^,…, *G^s^*, *s* ≦ 0, be the minimal proper faces of *C*. Since 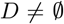 and 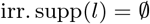, for all *l* ∈ *L*, we have *C* ≠ L. Thus *C* has at least one minimal proper face and *s* ≥ 1. By Prop. 2.2, 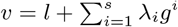, for some *l* ∈ *L* and λ_*i*_ ≥ 0, *g^i^* ∈ *G^i^* \ *L*, for *i* = 1,…, *s*. It follows irr. 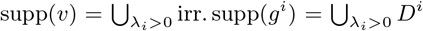, where *D^i^* = irr. supp(*g^i^*) is the MMB of the minimal proper face *G^i^*, for *i* = 1,…,*s*.

Next we generalize the characterization of minimal proper faces by MMBs [10] to higher-dimensional faces.

#### Proposition 4.2.

*Let C be the flux cone of a metabolic network and let F be a face of C. Then all v* ∈ relint(*F*) *have the same irreversible support or equivalently share the same metabolic behavior.*

*Proof.* Let *v,w* ∈ relint(*F*). Assume w.l.o.g. that there exists *j* ∈ irr. supp(*w*) \ irr. supp(*v*). Then *v_j_* = 0, but *w_j_* > 0, and hence for any λ > 1, we have λ*v* + (1 – λ)*w* ∈ *C*. However, since *v,w* ∈ relint(*F*), we know λ*v* + (1 – λ)*w* ∈ *F*, for some λ > 1. This shows irr. supp(*v*) = irr. supp(*w*), which implies the statement.

#### Example 4.3.

*Consider the network in Fig. 1. If we remove the redundant irreversibility constraint v*_6_ ≥ 0 *and assume* 6 ∈ Rev, *the MMBs of the network are* {2}, {7}, {8} (*if* 6 ∈ Irr, *the MMBs are* {2}, {6, 7}, {6, 8}*). The face lattice together with the EFMs contained in each face is shown in Fig. 2.*

**Figure 2:**
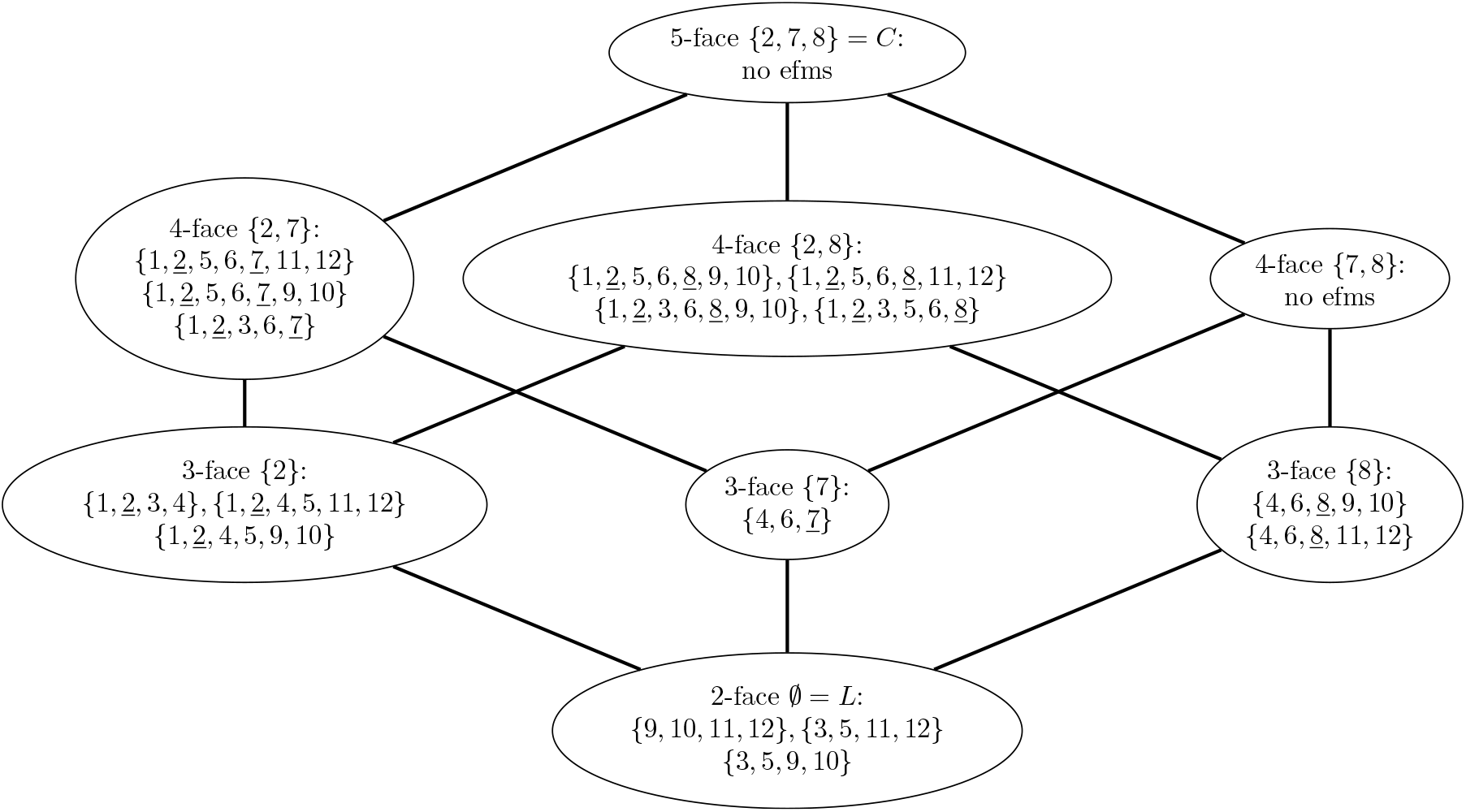
Face lattice of the network in Fig. 1. Each node represents a face of the flux cone together with the corresponding metabolic behavior and the EFMs contained in that face. The active irreversible reactions are underlined. The only 2-face is the lineality space *L* and the only 5-face is the entire flux cone *C*

## 5 The degree of flux vectors

Let *C* be the flux cone of a metabolic network. We define the *degree* deg(*v*) of a flux vector *v* ∈ *C* as the dimension of the inclusionwise minimal face of *C* containing *v*, which for *v* ≠ 0 is the unique face *F* of *C* with *v* ∈ relint(*F*). By Prop. 2.1, we have

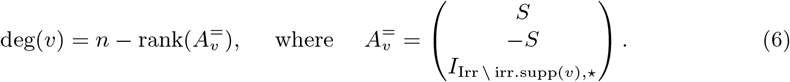

It follows that flux vectors in the lineality space *L* of *C* have degree dim(*L*) and flux vectors in minimal proper faces have degree dim(*L*) + 1. A flux vector in the relative interior of *C* has degree dim(*C*). Next we further characterize these vectors.

### Proposition 5.1.

*Let C* = {*v* ∈ ℝ^*n*^ | *Sv* = 0, *v*_Irr_ ≥ 0} *be a flux cone with no implicit equalities in v*_Irr_ ≥ 0. *For v* ∈ *C we have* deg(*v*) = dim(*C*) *if and only if v_i_* > 0 *for all i* ∈ Irr.

*Proof.* Direct consequence of Prop. 2.4.

Although small examples with EFMs in the relative interior of the flux cone can be constructed, we note that real biological networks typically do not have EFMs where all unblocked irreversible reactions are active. In these cases, there are no EFMs in the relative interior of the cone, i.e., all EFMs lie on the relative boundary of *C*.

Next we prove an upper bound on the degree of flux vectors.

#### Proposition 5.2.

*Let C* = {*v* ∈ ℝ^*n*^ | *Sv* = 0, *v*_Irr_ ≥ 0} *be the flux cone of a metabolic network with lineality space L. Then for each flux vector v ∈ C*

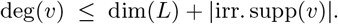

*Proof.* By definition of the lineality space *L, t* := dim(*L*) = *n* – rank(*A*), with 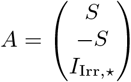.

By (6), we have 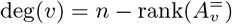, with 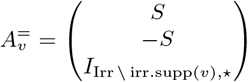.

It follows 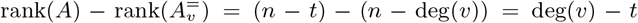. Thus, at least deg(*v*) – *t* rows from *A* must be missing in 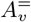. This implies |irr. supp(*v*)| ≥ deg(*v*) – *t* or deg(*v*) ≤ *t* + |irr. supp(*v*)|.

By combining Prop. 3.4 and Prop. 5.2, we get an upper bound on the degree of EFMs.

#### Corollary 5.3.

*Let C* = {*v* ∈ ℝ^*n*^ | *Sv* = 0,*v*_Irr_ ≥ 0} *be the flux cone of a metabolic network with lineality space L. Then for each EFM e* ∈ *C*

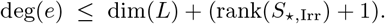

The following example shows that the bound from Cor. 5.3 is optimal.

#### Example 5.4.

*The network in Fig. 3 contains 2 metabolites and 4 reactions, with* Rev = {1, 2} *and* Irr = {3, 4}. *Given the stoichiometric matrix*

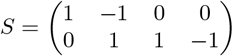

*the network has the EFMs e*^1^ = (1, 1, 0, 1), *e*^2^ = (–1, –1,1, 0) *and e*^3^ = (0, 0,1,1), *with* deg(*e*^1^) = deg(*e*^2^) = 1 *and* deg(*e*^3^) = 2. *Note that C* = cone({*e*^1^, *e*^2^}) *and e*^3^ = *e*^1^ + *e*^2^ ∈ relint(*C*). *Since there are no reversible flux vectors, we have* dim(lin. space(*C*)) = 0. *Furthermore*, rank(*S*_*,Irr_) = 1 *and thus* deg(*e*^3^) = dim(lin. space(*C*)) + (rank(*S*_*,Irr_) + 1) = 2.

**Figure 3:**
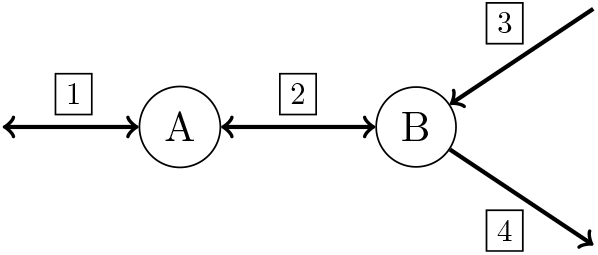
Example network

For the example networks `E.coli core, Pentose Phosphate Pathway` and `Pyruvate` from Sect. 3.3, the maximum degree of EFMs and the upper bounds from Prop. 5.2 and Cor. 5.3 are given in Tab. 2. We can see that *l* = max{|irr. supp(*e*)| : *e* EFM} is much smaller than the upper bound rank(*S*_*,Irr_) +1 from Prop. 3.4. The actual degrees of the EFMs in these networks are summarized in Fig. 4.

**Figure 4:**
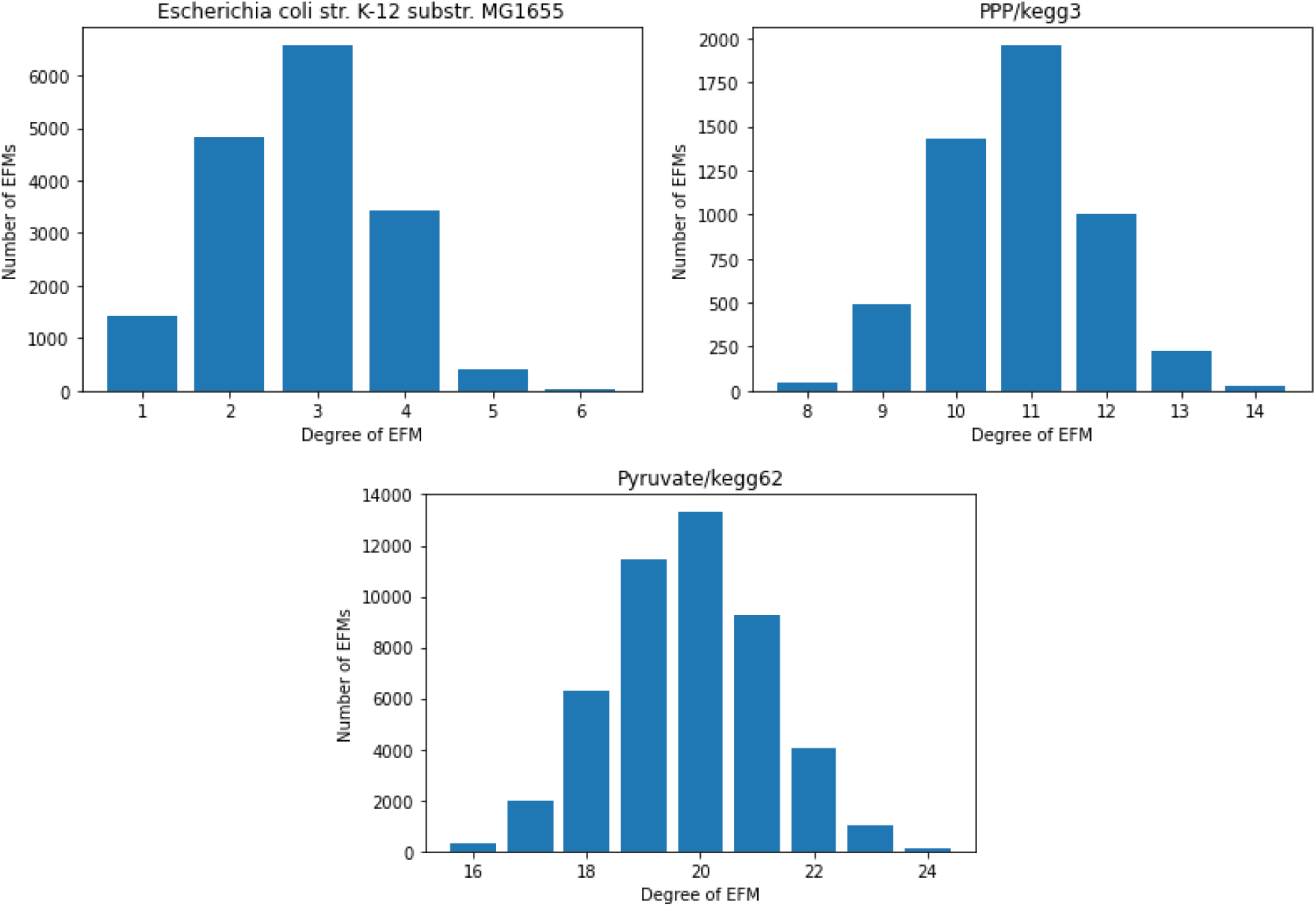
Degree distribution of EFMs

**Table 2:** Maximum number of active irreversible reactions and maximum degree of EFMs together with the upper bounds from Prop. 3.4, 5.2 and Cor. 5.3

|  | E. coli core | Pentose Phosphate | Pyruvate |
| --- | --- | --- | --- |
| $(\vert \mathcal{M}\vert , \vert \mathcal{R}\vert )$ | (72, 94) | (34, 57) | (28, 81) |
| $\vert \text{Irr}\vert $ | 48 | 19 | 40 |
| $\vert \text{Rev}\vert $ | 46 | 38 | 41 |
| $\text{rank}(S)$ | 67 | 34 | 28 |
| $\text{rank}(S_{*,\text{Irr}})$ | 41 | 16 | 24 |
| $t = \dim(\text{lin. space}(C))$ | 0 | 8 | 16 |
| $q = \max\{\vert \text{irr. supp}(e)\vert : e \text{ EFM}\}$ | 23 | 9 | 10 |
| $r = \text{rank}(S_{*,\text{Irr}}) + 1$ (cf. Prop. 3.4) | 42 | 17 | 25 |
| $\max\{\deg(e) : e \text{ EFM}\}$ | 6 | 14 | 24 |
| $t + q$ (cf. Prop. 5.2) | 23 | 17 | 26 |
| $t + r$ (cf. Cor. 5.3) | 42 | 25 | 41 |
| $\dim(C)$ | 23 | 23 | 53 |
| $\vert \text{EFMs}\vert $ | 16673 | 5180 | 47854 |

The next proposition explains the scarcity of EFMs in the relative interior of a flux cone *C*, in the facets of *C* and in the faces of dimension dim(*C*) – 2.

#### Proposition 5.5.

*Let C* = {*v* ∈ ℝ^*n*^ | *Sv* = 0,*v*_Irr_ ≥ 0} *be a flux cone with no redundant inequalities or implicit equalities in v*_Irr_ ≥ 0. *If* |Irr| > rank(*S*_*,Irr_) + *q, for some q* ∈ {1, 2, 3}, *then* deg(*e*) ≤ dim(*C*) – *q, for each EFM e of C*.

*Proof.* By Prop. 3.4, |irr. supp(*e*)| ≤ rank(*S*_*,Irr_) + 1, for each EFM e of *C*.

Assume deg(*e*) = dim(*C*) – (*q* – 1), for some *q* ∈ {1, 2, 3}. Then, by definition, the inclusionwise minimal face *F* containing e has dimension dim(*C*) – (*q* – 1) ≥ dim(*C*) – 2. It follows that *e* resp. *F* is contained in exactly *q* – 1 facets of *C*. Here we use for *q* = 3 that a (dim(*C*) – 2)-face is contained in exactly two facets, cf. [14, p.105].

By the hypothesis on the description of the cone, it follows |irr. supp(*e*)| = |Irr| – (*q* – 1). So we get |Irr| – (*q* – 1) ≤ rank(*S*_*,Irr_) +1 or |Irr| ≤ rank(*S*_*,Irr_) + *q*, in contradiction to the hypothesis |Irr| > rank(*S*_*,Irr_) + *q*.

In the proof we used that (dim(*C*) – *q*)-faces of a cone *C* are contained in exactly *q* facets of *C*, for *q* = 0, 1, 2. As the example of a 3-dimensional pointed cone with *n* facets shows, a (dim(*C*) – 3)-face (here the origin) can be contained in an arbitrary number of facets, and thus a similar argument does not hold for such faces. To limit the number of facets a face can be contained in, we introduce the concept of *l-simple* cones and use this for a generalization of Prop. 5.5.

A cone *C* ⊆ ℝ^*n*^ is called *l-simple* for some *l* ≥ 1, if every *k*-face of *C* is contained in at most *l* · (dim(*C*) – *k*) facets of *C*, for all *k* = dim(lin. space(*C*)),…, dim(*C*). Assuming that a flux cone is *l*-simple leads to another bound on the degree of EFMs.

#### Proposition 5.6.

*Let C* = {*v* ∈ ℝ^*n*^ | *Sv* = 0,*v*_Irr_ ≥ 0} *be an l-simple cone with no redundant inequalities or implicit equalities in v*_Irr_ ≥ 0. *Then for each EFM e* ∈ *C*

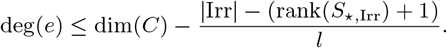

*Proof.* By Prop. 3.4, |irr. supp(*e*)| ≤ rank(*S*_*,Irr_) + 1 for each EFM *e* ∈ *C* and thus at least |Irr| – (rank(*S*_*,Irr_) + 1) entries of *v*_Irr_ are equal to zero. Hence *e* is contained in at least |Irr| – (rank(*S*_*,Irr_) + 1) facets of *C*. Suppose deg(*e*) = *k* and let *e* ∈ *F*, where *F* is a *k*-face of *C*. Since *C* is *l*-simple, *F* is contained in at most *l* · (dim(*C*) – *k*) facets. It follows

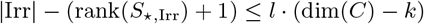

or

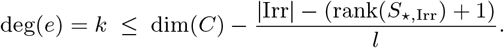

Note that this bound is mainly theoretical because for the computation of l all faces of the flux cone have to be considered. Nevertheless |Irr| ≦ rank(*S*_*,Irr_) and |Irr| is typically significantly larger than rank(*S*_*,Irr_) (cf. Tab. 2).

## 6 The cardinality of minimal metabolic behaviors

We next prove an upper bound on the cardinality of MMBs.

#### Proposition 6.1.

*Let C be the flux cone of a metabolic network* 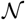 *with lineality space L. Then for each MMB D*

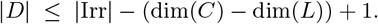

*Proof.* By definition of an MMB, there exist a minimal proper face *G* of *C*, dim(*G*) = dim(*L*) + 1, and a flux vector *g* ∈ *G* \ *L* such that *D* = irr. supp(*g*). It follows that *g* is contained in at least dim(*C*) – (dim(*L*) + 1) facets of *C*. Therefore at least dim(*C*) – (dim(*L*) + 1) inequalities in *v*_Irr_ ≥ 0 are satisfied by *g* with equality, which implies |irr. supp(*g*)| = |*D*| ≤ |Irr| – (dim(*C*) – (dim(*L*) + 1)).

In general, MMBs often contain irreversible reactions for which the non-negativity constraint is redundant. If we remove redundant non-negativity constraints (i.e., shift the corresponding reactions from Irr to Rev) until the description contains no more redundant inequalities, this typically leads to much smaller cardinalities of MMBs.

For our example networks `E.coli core, Pentose Phosphate Pathway` and `Pyruvate`, the number of MMBs are 1421, 19 and 37 respectively. In Fig. 5 we compare the cardinalities of the MMBs in the original description of the flux cone and after removing redundant irreversibility constraints. If we start from the redundant description, Prop. 6.1 provides the upper bounds 18, 5 and 4 respectively, while the actual maximal sizes of the MMBs are 17, 4 and 3. If we remove redundant non-negativity constraints in lexicographical order (i.e., the redundant non-negativity constraint corresponding to the irreversible reaction with the smallest index is removed first), the bounds become sharp, i.e., we get the bounds 9, 3 and 1 respectively, and these bounds coincide with the actual maximal sizes of the MMBs.

#### Proposition 6.2.

*Let C* = {*v* ∈ ℝ^*n*^ | *Sv* = 0, *v*_Irr_ ≥ 0} *be the flux cone of a metabolic network with lineality space L and no redundant inequalities or implicit equalities in v*Irr ≥ 0. *Then the number of facets of C is equal to* dim(*C*) – dim(*L*) *if and only if each MMB has cardinality one*.

*Proof.* By Prop. 3.3, the number of facets is equal to |Irr|. Thus if |Irr| = dim(*C*) – dim(*L*), then by Prop. 6.1, 1 ≤ |*D*| ≤ |Irr| – (dim(*C*) – dim(*L*)) + 1 = 1, for each MMB *D* in *C*.

Conversely, if each MMB has cardinality 1, then each minimal proper face is the intersection of all facets of *C* but one. For all *i* ∈ Irr, *G^i^* = {*v* ∈ ℝ^*n*^ | *Sv* = 0,*v*_Irr_\{*i*} = 0,*v_i_* ≦ 0} is a face of *C*, with dim(*G^i^*) ≤ dim(*L*) + 1. Since *v_i_* ≥ 0 is not an implicit equality, there exists *g^i^* ∈ *C* with 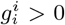. Let 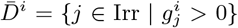 be the metabolic behavior defined by *g^i^*. By Prop. 4.1, 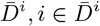, is the union of MMBs, which by hypothesis all have cardinality 1. Thus, for all *i* ∈ Irr, *D^i^* = {*i*} is an MMB with the corresponding minimal proper face *G^i^*, where *G^i^* \ *L* = {*v* ∈ ℝ^*n*^ | *Sv* = 0, *v*_Irr_\{*i*} = 0, *v_i_* > 0} and dim(*G^i^*) = dim(*L*) + 1. We conclude that the number of minimal proper faces of *C* is equal to the number of facets, which by Prop. 3.3 is equal to | Irr|.

It remains to prove that |Irr| = dim(*C*) – dim(*L*). Let *U* = {*u* ∈ ℝ^*n*^ | *Su* = 0} and *W* = {*w* ∈ ℝ^*n*^ | *w*_Irr_ = 0}. Then *U* ⋂ *W* = *L* and since by hypothesis there are no implicit equalities, dim(*C*) = dim(aff(*C*)) = dim(*U*). From the dimension formula, we get dim(*U*+*W*) = dim(*U*) + dim(*W*) – dim(*U* ⋂ *W*) = dim(*C*) + (*n* – |Irr|) – dim(*L*) or dim(*C*) – dim(*L*) = |Irr| – (*n* – dim(*U* + *W*)).

We claim dim(*U* + *W*) = n, i.e., *U* + *W* = ℝ^*n*^. For each minimal proper face *G^i^,i* ∈ Irr, choose *e^i^* ∈ *G^i^* \ *L* with 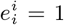. Then 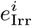 is a unit vector, for all *i* ∈ Irr. Given *v* ∈ ℝ^*n*^, let 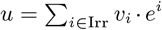 and 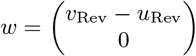. Since *Se^i^* = 0, *i* ∈ Irr, we get *Su* = ∑_*i*∈Irr_ *v_i_* · *Se^i^* = 0 and thus *u* ∈ *U*. By definition, *w* ∈ *W*. For all *j* ∈ Irr, we have 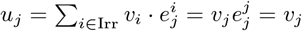, and thus *u*_Irr_ = *v*_Irr_. Altogether, we get 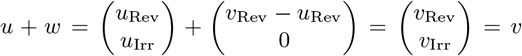, which shows *U* + *W* = ℝ^*n*^.

**Figure 5:**
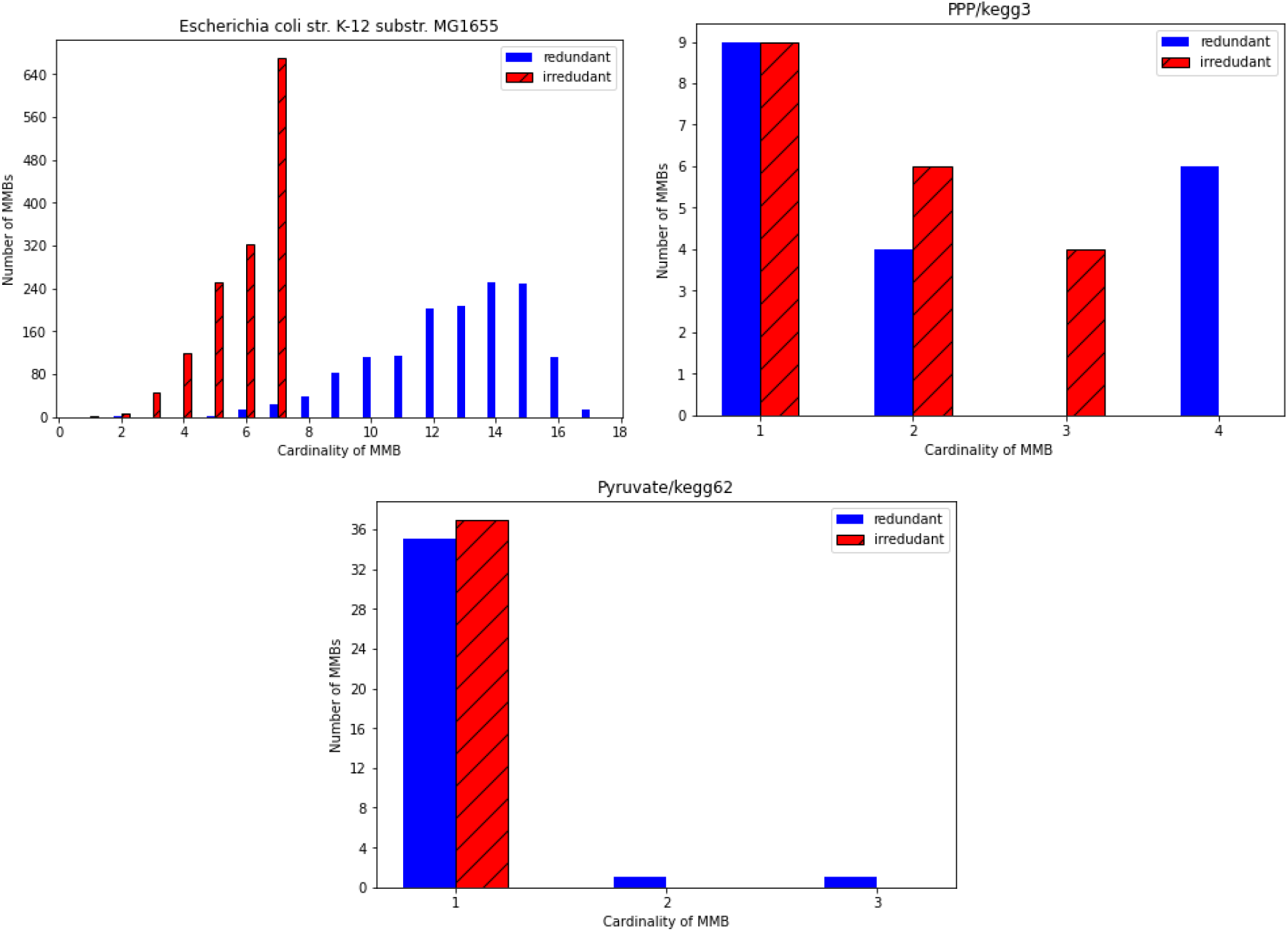
Cardinalities of MMBs with and without redundant irrversibility constraints

## 7 Conclusion

In this paper, we studied geometric properties of elementary flux modes in metabolic networks. To structure the set of EFMs, we introduced the degree of an EFM. We derived upper bounds on the degree and studied the distribution of EFMs on the face lattice of the flux cone. In our future work, we plan to build on these geometric insights to develop new algorithmic approaches for elementary mode computation.

## Acknowledgments

FW was partially supported by project AA3-11* of MATH+, the Berlin Mathematics Research Center.

## Data Availability

Data was obtained from public domain resources as cited in the paper.

